# PlotTwist – a web app for plotting and annotating time-series data

**DOI:** 10.1101/745612

**Authors:** Joachim Goedhart

## Abstract

The results from time-dependent experiments are often used to generate plots that visualize how the data evolves over time. To simplify state-of-the-art data visualization and annotation of data from such experiments, an open source tool was created with R/shiny that does not require coding skills to operate. The freely available web app accepts wide (spreadsheet) and tidy data and offers a range of options to normalize the data. The data from individual objects can be shown in three different ways: (i) lines with unique colors, (ii) small multiples and (iii) heatmap-style display. Next to this, the mean can be displayed with a 95% confidence interval for the visual comparison of different conditions. Several color blind friendly palettes are available to label the data and/or statistics. The plots can be annotated with graphical features and/or text to indicate any perturbations that were applied during the time-lapse experiments. All user-defined settings can be stored for reproducibility of the data visualization. The app is dubbed PlotTwist and is available online: https://huygens.science.uva.nl/PlotTwist

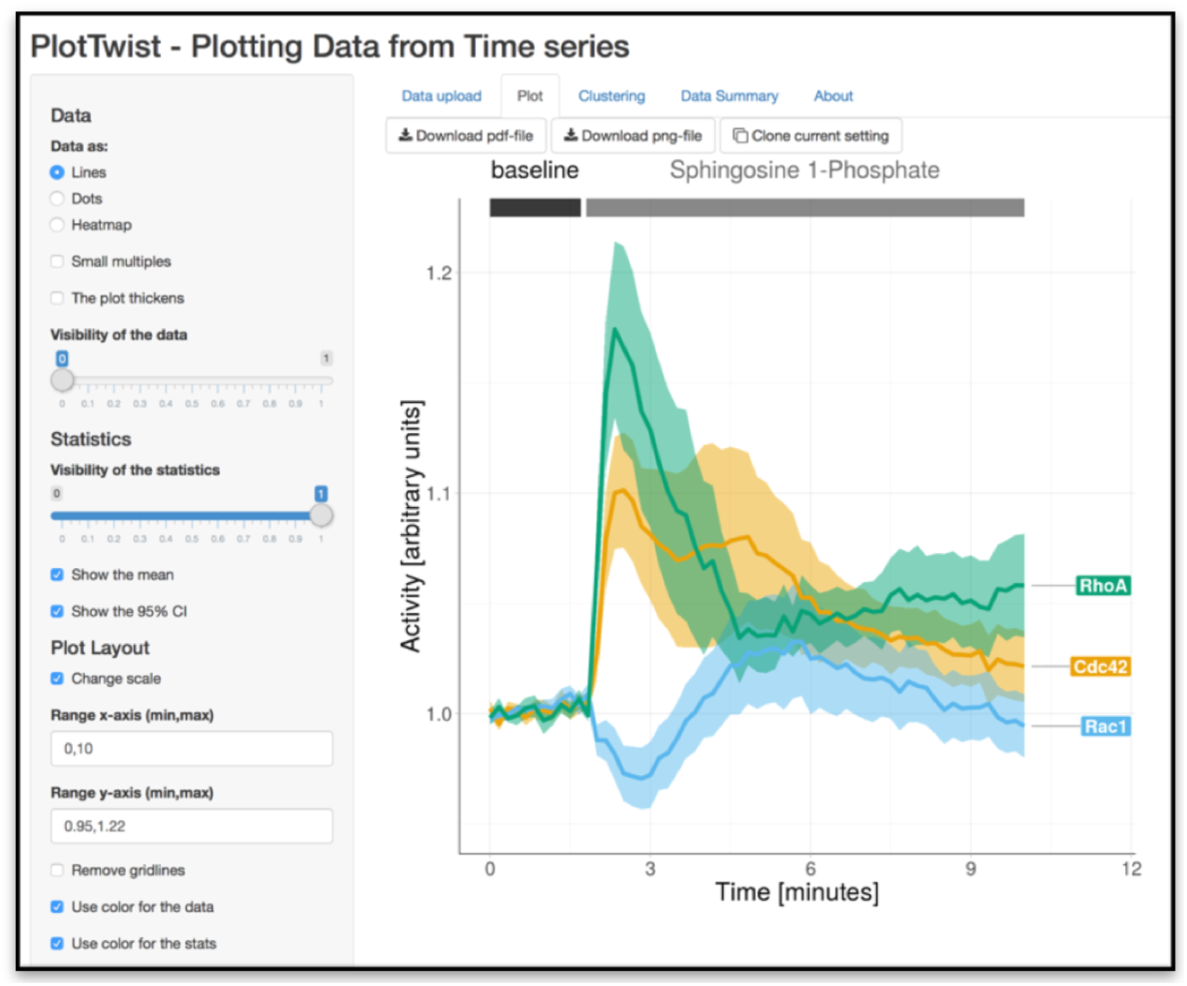

## Introduction

It has been stressed over the years that showing the actual data from individual measurements, instead of summaries only, gives a better picture of the actual data summaries (Weissgerber et al., 2015; Rousselet et al., 2017; Drummond and Vowler, 2011). This holds true for data acquired under static conditions as well as for time-dependent data to study dynamics. For static conditions, dotplots are a good way to graphically depict the data. Several online data visualization tools have been reported that include the option to display the actual data (Weissgerber et al., 2017; Postma and Goedhart, 2019; Goedhart, 2019; Ho et al., 2019).

However, similar online tools for the visualization of continuous data seem (to the best of our knowledge) to be lacking. This is surprising, since understanding dynamic systems, especially relevant in biology, requires experimentation over time. In addition, time-series experiments can be high content (Megason and Fraser, 2007) and may include the application of multiple, different perturbations. The complexity of time-series data requires a tool to rapidly inspect the data and have access to different visualizations. Next to that, it is important to clearly communicate the treatments/perturbations that were applied during the experiment. Ideally, a data visualization tool for time-series data would be freely available, open source and generate high quality graphs that are publication ready. Having previously reported such a webtool for static data (Postma and Goedhart, 2019) and given the enthusiastic response by the community, we were motivated to generate a webtool for time-series data.

The new web-tool that we generated is dubbed PlotTwist, for plotting data from Time-lapse experiments With Indicators of conditions at Set Times. This free and open source app is developed to facilitate the rapid visualization and annotation of time-lapse data. Creating graphs with PlotTwist does not require coding skills, yet it produces the state-of-the-art data visualization offered by the ggplot2 package. Therefore, we think that PlotTwist is well-suited to inspect data and generate publication quality visualizations of data from time-lapse experiments. The features and use of PlotTwist are highlighted below.

## Availability, code & issues

The PlotTwist webtool is available at: https://huygens.science.uva.nl/PlotTwist or https://goedhart.shinyapps.io/PlotTwist/

The code was written in R using R (https://www.r-project.org) and Rstudio (https://www.rstudio.com). To run the app several freely available packages are required: shiny, ggplot2, dplyr, tidyr, readr, readxl, magrittr, ggrepel, DT, dtw (Giorgino, 2009). The code of the version 1.0.3 that is reported in this manuscript is archived at Zenodo.org: https://doi.org/10.5281/zenodo.3384426

Up-to-date code and new release will be made available on Github, together with information on running the app locally: https://github.com/JoachimGoedhart/PlotTwist

The Github page of PlotTwist is the preferred way to communicate issues and request features (https://github.com/JoachimGoedhart/PlotTwist/issues). Alternatively, the users can contact the developers by email or <u>Twitter</u>. Contact information is found on the “About” page of the app.

## Data input, structure and normalization

The data can be provided by copy-paste or via upload of a CSV or XLS file. The data structure can be either a wide, spreadsheet type format or a long, tidy format. Example data, of which the details can be found in earlier publications (Reinhard et al., 2017; Mastop et al., 2017) is included in the app.

### Wide data format

For the wide format, the first column is assumed to contain the time-points. Each of the other columns is taken as data of different samples. The wide format will be converted into the tidy format (Figure 1). This conversion is not visible for the user, but it is required for plotting with ggplot2. When multiple wide datasets are simultaneously uploaded (in CSV or XLS format) each of these is treated as a different condition. The supplemental movie S1 demonstrates how the multiple file upload is done. The filename is used as a label for the condition.

**Figure 1:**
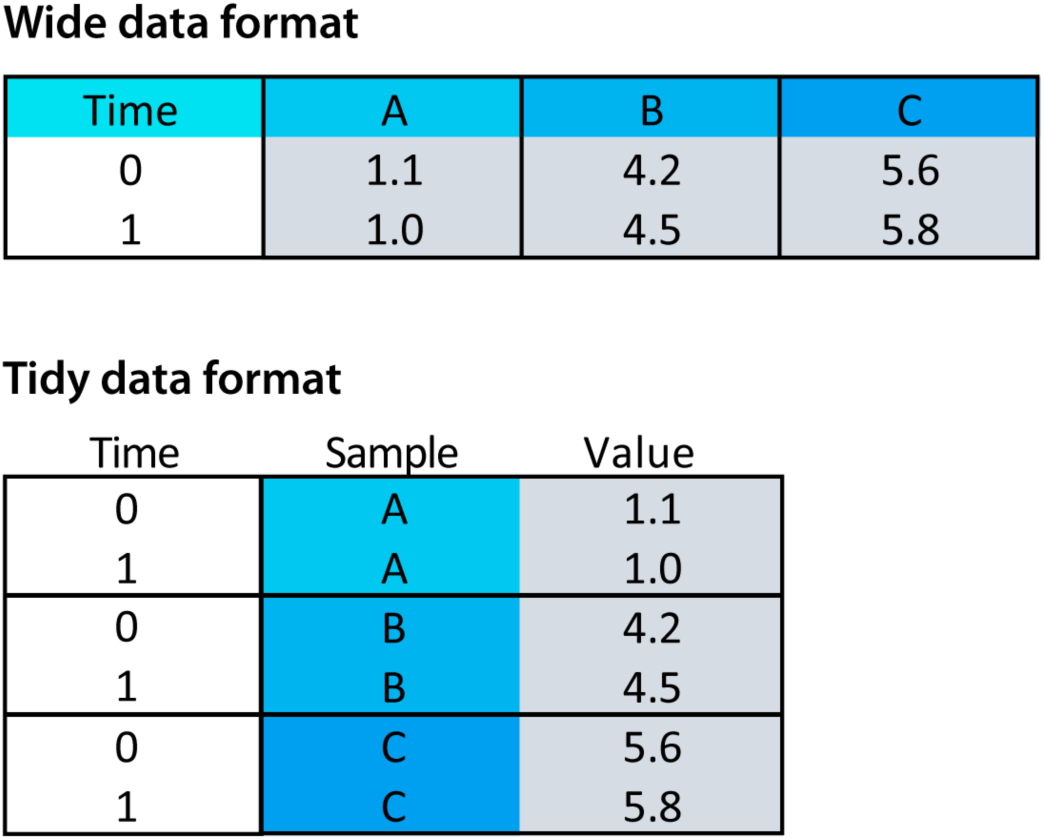
The wide versus tidy data format for time dependent data. In the wide (spreadsheet-like) data format the first column holds the time data and the other columns represent data from different objects (A,B & C). In the tidy format each column is a single variable and each row is an observation. Note that the time data is repeated for each of the objects A,B & C.

### Tidy data format

For data in a tidy data format, the user is asked to select the columns that have (i) the time data, (ii) the measured data (values) (iii) a unique identifier for each sample and (iv) the condition.

After data upload, there are several options to normalize the data, to facilitate comparison. Some of the normalization options are: divide values over a baseline value (I/I0), divide a difference by the baseline (ΔI/I0), divide by the maximum value and scale between 0 and 1. The normalized data in a tidy format can be downloaded as a CSV file (comma separated values).

Finally, there is an option to deselect data from individual objects.

## Data visualization

The default graph shows time along the x-axis and the measuremed values on the y-axis. A line plot is constructed by connecting all the coordinates from each sample by lines. The mean value and/or the 95% confidence interval of the mean can be drawn as a transparent layer on top of the lines from individual samples. The 95% confidence intervals enable the comparison of difference conditions by ‘visual inference’ (Cumming and Finch, 2005; Cumming et al., 2007; Gardner and Altman, 1986). The line thickness and transparency of the individual data and statistical summary can be adjusted by the user. Several color palettes that are colorblindness friendly are available for the individual data and the statistics. When data for multiple conditions is provided, the statistics for each of the conditions can be shown with different colors. By tweaking the transparency, color and line thickness, the display of the individual data and summary can be optimized.

## Display of data from individual objects

When multiple objects (cells) are measured simultaneously, it is often useful to examine the responses of the individual objects. To this end, PlotTwist can show the data in three different ways: (i) a lineplot, where single time-traces can be distinguished by color and/or labels, (ii) small multiples showing each response in a separate graph and (iii) a heatmap-style representation that shows the response of each object in false color. The lineplot is effective when the number of objects is low to medium (up to 20). The small multiples may still work well for up to 100 objects. The heatmap-style representation is especially suited for the display of rich datasets with many objects (>10 objects up to several hundreds). Figure 2 shows the same data in an ordinary lineplot (panel A) and in a heatmap style presentation (panel B). When a heatmap is used, the data from the different objects can be sorted according to several criteria, including the maximal signal, integrated response and alphabetical order.

**Figure 2:**
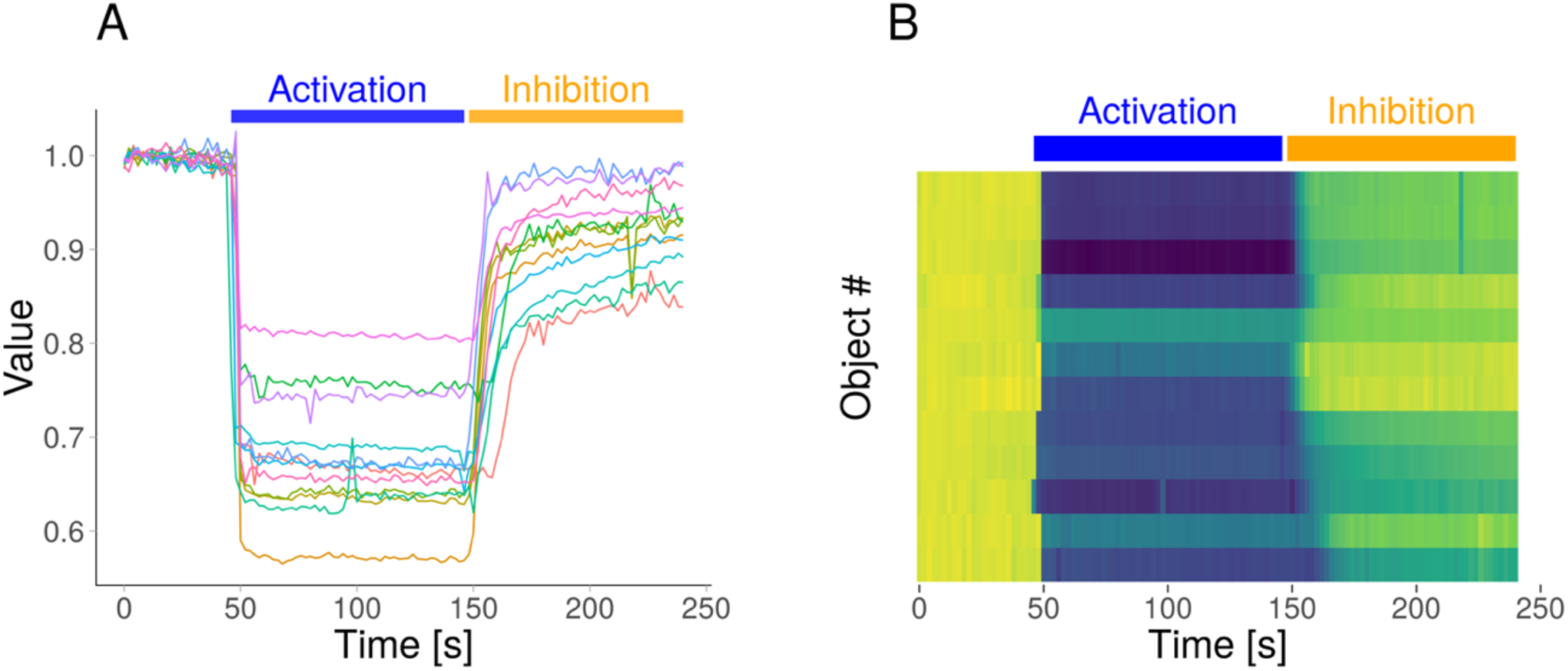
Different plots generated in PlotTwist with the same data. An ordinary lineplot, showing the individual data as lines (A) and a heatmap style presentation (B) of the (same) example data. Both graphs are annotated with bars on top of the plot to display an activating and an inhibiting perturbation.

## Color

Color is an important feature for distinguishing objects or conditions. To distinguish between different categories *qualitative* color schemes are used. In PlotTwist, several qualitative color schemes are implemented that are color blind friendly. The palettes have been designed by Masataka Okabe & Kei Ito (https://jfly.uni-koeln.de/color/) and Paul Tol (https://personal.sron.nl/~pault/). The palettes are composed of 7-10 colors and their exact composition is shown in figure 3A. The palette from Okabe and Ito is the default in PlotTwist. The most suitable color palette needs to be on a case-by-case basis. To give a rough idea on the performance of each of the palettes, we have used them to give unique colors to 6 different lines in a realistic plot (figure 3B).

**Figure 3:**
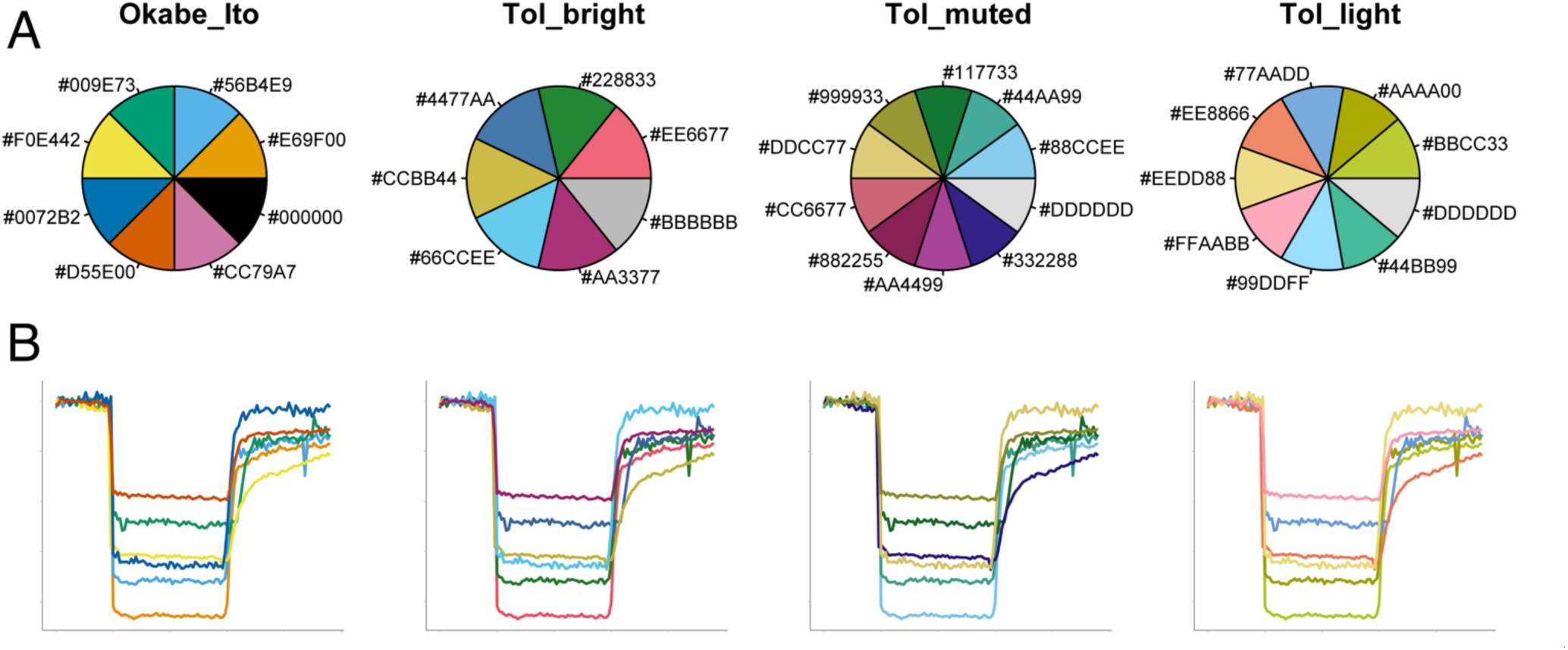
Composition and application of the color blind friendly palettes that are available in PlotTwist. (A) An overview of the color composition of the qualitative color palettes that were designed to have colors that are distinct to all people, including those with a color vision deficiency. The RGB code for each of the colors is indicated in hexadecimal code, preceded with a hashtag. (B) A realistic application of the color palettes shown in the upper panel for the unique labeling of 6 different lines.

Factors that affect the choice of the palette may be the shape and thickness of the lines (thicker lines are usually easier to identify), their position and overlap and the number of unique colors that is needed. In addition, the medium that is used to display the colors affects their appearance. The colors will look different on a screen, on print or when projected with a beamer. It may be worthwhile to ask person with a color vision deficiency which palettes work best, although color blindness is a heterogeneous deficiency (Frane, 2015).

Alternatively, user-defined colors can be applied by either typing the name or the hexadecimal color code. An example, also available in PlotTwist, of three different colors: “turquoise2, #FF2222, lawngreen”, where #FF2222 is a hexadecimal code for red. Whenever there are more objects than colors, the color scheme will be repeated, and therefore different objects will be labeled with the same color.

For the heatmap style presentation the perceptually uniform and colorblindness friendly viridis color palette is used (https://bids.github.io/colormap/).

## Labeling of data

Colors can be supplemented with, or replaced by, direct labeling of the data. To this end, PlotTwist offers an option to add labels to the objects on the right side of the plot. The ggrepel package is used to reduce overlap between the labels. The labels show the name of the data. When only data is shown (no statistics), each individual line is labeled. In the case that the average curve is displayed, the label indicates the condition, next to the thick line that shows the average. When colors are used, the label has the same color as the data. Labels have the advantage that they (i) have a relatively large area filled with color, (ii) indicate the condition with text in addition to color and (iii) are displayed next to the data. Therefore, labels can effectively replace legends. The use of labels instead of a standard legend is compared in figure 4A & 4B.

**Figure 4:**
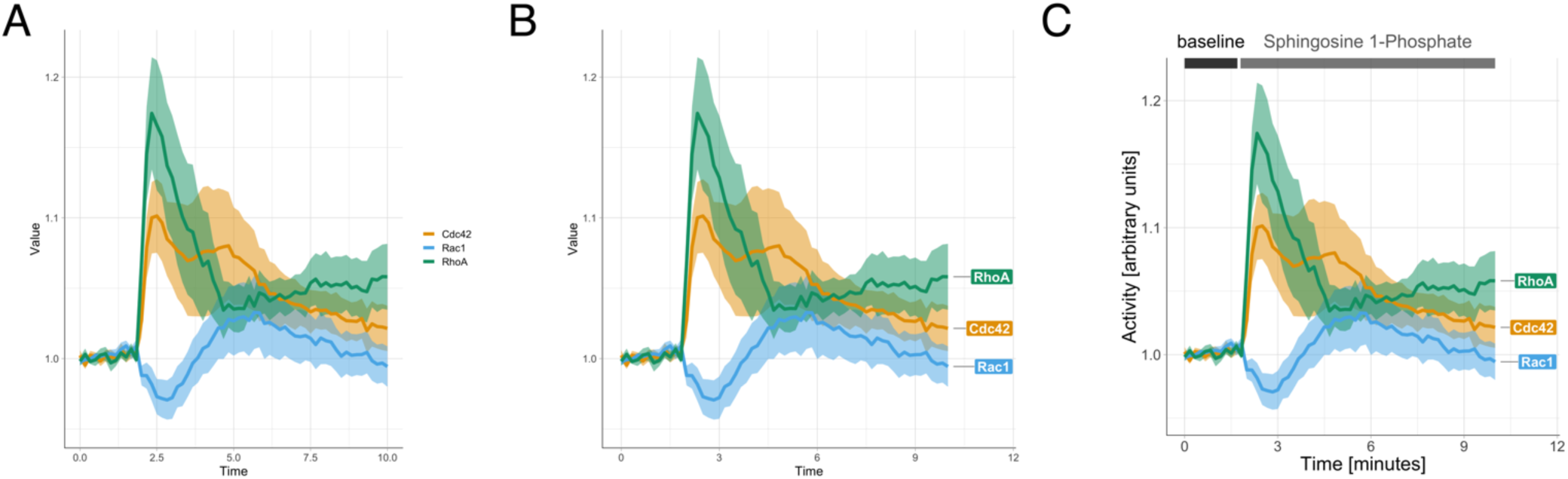
Different annotation styles that are available in PlotTwist. From left (A) to right (C), increasingly explicit labeling is used to explain the data. The data is the same for each plot and shows the relative activity of the proteins Cdc42, Rac1 and RhoA in response to treatment with sphingosine 1-phosphate (Reinhard et al., 2017). (A) A standard legend is used to explain the color used for the lines. (B) Direct annotation of the lines is used by placing a label next t the data. In addition, the color of the label is the same as the color of the data. (C) A fully annotated plot with clear labeling of the axes, direct annotation of the data and a visualization of the treatments at the different times.

## Indicating treatments

Time-lapse experiments are uniquely suited to analyze the effects of multiple treatments or perturbations during a single measurement. Some examples of perturbations in biological experiments are (i) the application of an agonist to stimulate signal transduction, (ii) the addition of a drug to inhibit proteins or stimulate dimerization or (iii) illumination of light sensitive proteins (optogenetics) to direct cellular processes. The annotation of plots to indicate the treatments is often done manually. This can be time-consuming and inaccurate. To address this issue, PlotTwist enables straightforward and reproducible annotation of an unlimited number of treatments. The annotation can be added as a bar above the plot, a box in the plot or a combination. The color of the annotation can be adjusted and text can be added. An example of annotations is shown in figure 2A&B and figure 4C, where the treatments are indicated by bars on top of the plot. Next to bars, there is also an option to use boxes or a combination of a bar and box.

## Output

The data that is used for the plot can be downloaded in a CSV format in the tidy format. Any deselected data will not appear in the file. In case that the data was subjected to normalization, there is also an option to download these data as a CSV file in tidy format. Downloaded CSV files can be used again by PlotTwist to plot the data.

The plot that is generated by the app can be directly retrieved by drag-and-drop from the web browser. In addition, the plot can be downloaded as a PNG or PDF file. The PNG is a lossless bitmap format. The PDF allows for downstream processing/editing with software that can handle vector-based graphics.

## Reproducibility

In general, graphical user interfaces offer substantial flexibility in the options and parameters that can be set. In PlotTwist, the user has control over scaling, annotation, color and more. Due to this flexibility, it is challenging to exactly reproduce the settings. Here, to achieve reproducibility, we have included an option to ‘clone’ the active setting. This action will generate a URL (Uniform Resource Locator) with all relevant parameters from the user interface. Through the URL, the parameters are fed back into the webtool and the settings are restored. In supplemental text S1, all the parameters (and their accepted values) that can be set through the URL are listed. As an example, figure 2A can be reproduced by employing this URL: https://huygens.science.uva.nl/PlotTwist/?data=1;;;fold;1,5;&vis=dataasline;1;;;1;&layout=%20;TRUE;;%20;;TRUE;;1;X;480;480&color=none&label=TRUE;A;TRUE;Time%20[s];Value;TRUE;30;24;18;8;;&sti m=TRUE;bar;46,146,148,240;Activation,Inhibition;Blue,Orange&

The plot in figure 4C can be recreated by using this link: https://huygens.science.uva.nl/PlotTwist/?data=2;TRUE;;fold;1,5;&vis=dataasline;0;TRUE;TRUE;1;;&la yout=;;TRUE;;0.95,1.22;TRUE;TRUE;6;X;600;600&color=none&label=;;TRUE;Time [minutes];Activity [arbitrary units];TRUE;24;24;18;8;;;TRUE&stim=TRUE;bar;0,1.7,1.

Moreover, the option to set parameters through a URL allows to make presets. For instance, when a certain set of graphs need to be annotated in the same way this can be achieved by passing the relevant parameters through the URL. For annotation with three labels (baseline, activation, inhibition) at times 10-40,60-140,170-240, the following URL can be used: https://huygens.science.uva.nl/PlotTwist/?stim=TRUE;bar;10,40,60,140,170,240;baseline,activation,inhibition

After using the URL to start PlotTwist, the data can be uploaded and the (preset) annotation will be present in the plot that visualizes the imported data.

## Conclusion

Performing measurements over time on several objects is an important experimental strategy. The resulting data can be high-content (when many objects are measured) and complex (due to application of different treatments or perturbations). To evaluate the results, data visualization tools that can rapidly switch between different presentation modes are needed. To communicate the results, state-of-the-art data visualization tools that enable clear annotations are needed. We have developed the freely available and open source tool PlotTwist, to serve both purposes. It can be used for inspection of data and rapidly switch between different presentations, i.e. standard line plot, small multiples and heatmap. Basic statistics (mean and 95%CI) can be added in user defined transparency and color. To generate publication-ready figures we use the state-of-the-art graphical package (ggplot2) to make the plots. The flexible and reproducible annotation of both data and experimental conditions allows for clear communication of the results. In summary, PlotTwist allows anyone to inspect data and to generate state-of-the art data visualizations. Although this application was written with biological processes in mind, it could be of used for any dataset from time-dependent measurements.

## Acknowledgments

Some of the PlotTwist code is taken from PlotsofData and it is partially inspired by ggplotGUI (https://github.com/gertstulp/ggplotgui/) by Gert Stulp. The colorblind safe palettes were developed by Paul Tol (https://personal.sron.nl/~pault/) and Masataka Okabe & Kei Ito (https://jfly.uni-koeln.de/color/). We are grateful to our colleagues at Molecular Cytology (UvA) for helpful discussions and Auke Folkerts (UvA, The Netherlands) for help with the server that runs shiny. Finally, the feedback, suggestions, enthusiastic responses and example plots that are shared on Twitter are highly appreciated.

