## Supplemental Text S1 for "PlotTwist – a web app for plotting and annotating time-series data"

**S1 Text – Passing parameters to PlotTwist through the HTML address**

**Background information on setting parameters through the HTML address**

Passing parameters through the HTML address can be used to define a visualization and alter the standard layout. There are several queries (?data, ?vis, ..) that can be used. Each of the queries can hold multiple parameters. The queries are separated by an ampersand (&) and the parameters are separated by a semicolon(;).

Parameters that need no change (i.e. remain default) should be left empty. The parameters are defined by their position so their order is critical. Below, the parameters than can be changed through the HTML address are indicated, with the options between brackets:

?data

[1] data_input	{1/2/3/4/5}

[2] tidyInput	{T}

[3] normalization	{T}

[4] norm_type	{max/min/zero_one/integral/diff/z-score/perc/fold}

[5] base_range	-> number (minimum value),number (maximum value)

?vis

[1] data_form	{dataasline/dataasdot|dataaspixel}

[2] alphaInput	{0-1}

[3] summaryInput	{median/mean/box/violin}

[4] add_CI	{T}

[5] alphaInput_summ	{0-1}

[6] multiples	{T}

[7] thicken	{T}

?layout

[1]	{T}

[2] no_grid	{T}

[3] change_scale	{T}

[4] range_x	-> number (minimum value),number (maximum value)

[5] range_y	-> number (minimum value),number (maximum value)

[6] color_data	{T}

[7] color_stats	{T}

[8] adjustcolors	{1/2/3/4/5}

[9]

[10] plot_height	-> number reflecting pixels

[11] plot_width	-> number reflecting pixels

?color

[1] colour_list	{Condition}

[2] user_color_list	-> list of colors separated by comma’s

?label

[1] add_title	{T}

[2] title	-> text

[3] label_axes	{T}

[4] lab_x	-> text

[5] lab_y	-> text

[6] adj_fnt_sz	{T}

[7] fnt_sz_title	-> number

[8] fnt_sz_labs	-> number

[9] fnt_sz_ax	-> number

[10] fnt_sz_stim	-> number

[11] add_legend	{T}

[12] legend_title	-> text

[13] show_labels_y	{T}

?stim

[1] indicate_stim	{T}

[2] stim_shape	{bar/box/both}

[3] stim_range	list of numbers separated by comma’s

[4] stim_text	list of words separated by comma’s

[5] stim_colors	list of words separated by comma’s

Example URLs:

https://huygens.science.uva.nl/PlotTwist/?data=2;TRUE;;fold;1,5;&vis=dataasline;0.3;TRUE;TRUE;1;&layout= ;;; ;;;TRUE;6;X;480;600&color=none&label=TRUE;Rho GTPase;TRUE;Time;Activity;;24;24;18;8;TRUE;Condition&stim=TRUE;both;1.8,10;S1P;blue&

https://huygens.science.uva.nl/PlotTwist/?data=1;;;fold;1,5;&vis=dataasline;0.3;TRUE;TRUE;1;&layout= ;;; ;;;;1;X;480;600&color=none&label=;;TRUE;Time;;TRUE;24;24;18;8;;&stim=TRUE;bar;46,146,149,240;Histamine, Pyrilamine;black,blue&
